# Large-scale, high-resolution comparison of the core visual object recognition behavior of humans, monkeys, and state-of-the-art deep artificial neural networks

**DOI:** 10.1101/240614

**Authors:** Rishi Rajalingham, Elias B. Issa, Pouya Bashivan, Kohitij Kar, Kailyn Schmidt, James J. DiCarlo

## Abstract

Primates—including humans—can typically recognize objects in visual images at a glance even in the face of naturally occurring identity-preserving image transformations (e.g. changes in viewpoint). A primary neuroscience goal is to uncover neuron-level mechanistic models that quantitatively explain this behavior by predicting primate performance for each and every image. Here, we applied this stringent behavioral prediction test to the leading mechanistic models of primate vision (specifically, deep, convolutional, artificial neural networks; ANNs) by directly comparing their behavioral signatures against those of humans and rhesus macaque monkeys. Using high-throughput data collection systems for human and monkey psychophysics, we collected over one million behavioral trials for 2400 images over 276 binary object discrimination tasks. Consistent with previous work, we observed that state-of-the-art deep, feed-forward convolutional ANNs trained for visual categorization (termed DCNN_IC_ models) accurately predicted primate patterns of object-level confusion. However, when we examined behavioral performance for individual images within each object discrimination task, we found that all tested DCNN_IC_ models were significantly non-predictive of primate performance, and that this prediction failure was not accounted for by simple image attributes, nor rescued by simple model modifications. These results show that current DCNN_IC_ models cannot account for the image-level behavioral patterns of primates, and that new ANN models are needed to more precisely capture the neural mechanisms underlying primate object vision. To this end, large-scale, high-resolution primate behavioral benchmarks—such as those obtained here—could serve as direct guides for discovering such models.

**SIGNIFICANCE STATEMENT:** Recently, specific feed-forward deep convolutional artificial neural networks (ANNs) models have dramatically advanced our quantitative understanding of the neural mechanisms underlying primate core object recognition. In this work, we tested the limits of those ANNs by systematically comparing the behavioral responses of these models with the behavioral responses of humans and monkeys, at the resolution of individual images. Using these high-resolution metrics, we found that all tested ANN models significantly diverged from primate behavior. Going forward, these high-resolution, large-scale primate behavioral benchmarks could serve as direct guides for discovering better ANN models of the primate visual system.

## INTRODUCTION

Primates—both human and non-human—can typically recognize objects in visual images at a glance, even in the face of naturally occurring identity-preserving transformations such as changes in viewpoint. This view-invariant visual object recognition ability is thought to be supported primarily by the primate ventral visual stream (DiCarlo et al., 2012). A primary neuroscience goal is to construct computational models that quantitatively explain the neural mechanisms underlying this ability. That is, our goal is to discover artificial neural networks (ANNs) that accurately predict neuronal firing rate responses at all levels of the ventral stream and its behavioral output. To this end, specific models within a large family of deep, convolutional neural networks (DCNNs), optimized by supervised training on large-scale category-labeled image-sets (ImageNet) to match human-level categorization performance (Krizhevsky et al., 2012; LeCun et al., 2015), have been put forth as the leading ANN models of the ventral stream (Yamins and DiCarlo, 2016). We refer to this sub-family as DCNN_IC_ models (IC to denote ImageNet-categorization pre-training), so as to distinguish them from all possible models in the DCNN family, and more broadly, from the super-family of all ANNs. To date, it has been shown that DCNN_IC_ models display internal feature representations similar to neuronal representations along the primate ventral visual stream (Yamins et al., 2013; Cadieu et al., 2014; Khaligh-Razavi and Kriegeskorte, 2014; Yamins et al., 2014), and they exhibit behavioral patterns similar to the behavioral patterns of pairwise object confusions of primates (Rajalingham et al., 2015). Thus, DCNN_IC_ models may provide a quantitative account of the neural mechanisms underlying primate core object recognition behavior.

However, several studies have shown that DCNN_IC_ models can diverge drastically from humans in object recognition behavior, especially with regards to particular images optimized to be adversarial to these networks (Goodfellow et al., 2014; Nguyen et al., 2015). Related work has shown that specific image distortions are disproportionately challenging to current DCNNs, as compared to humans (RichardWebster et al., 2016; Dodge and Karam, 2017; Geirhos et al., 2017; Hosseini et al., 2017). Such image-specific failures of the current ANN models would likely not be captured by “object-level” behavioral metrics (e.g. the pattern of pairwise object confusions mentioned above) that are computed by pooling over hundreds of images and thus are not sensitive to variation in difficulty across images of the same object. To overcome this limitation of prior work, we here aimed to use scalable behavioral testing methods to precisely characterize primate behavior at the resolution of individual images and to directly compare leading DCNN models to primates over the domain of core object recognition behavior at this high resolution.

We focused on *core invariant object recognition*—the ability to identify objects in visual images in the central visual field during a single, natural viewing fixation (DiCarlo et al., 2012). We further restricted our behavioral domain to *basic-level* object discriminations, as defined previously (Rosch et al., 1976). Within this domain, we collected large-scale, high-resolution measurements of human and monkey behavior (over a million behavioral trials) using high-throughput psychophysical techniques—including a novel home-cage behavioral system for monkeys. These data enabled us to systematically compare all systems at progressively higher resolution. At lower resolutions, we replicated previous findings that humans, monkeys, and DCNN_IC_ models all share a common pattern of object-level confusion (Rajalingham et al., 2015). However, at the higher resolution of individual images, we found that the behavior of all tested DCNN_IC_ models was significantly different from human and monkey behavior, and this model prediction failure could not be easily rescued by simple model modifications. These results show that current DCNN_IC_ models do not fully account for the image-level behavioral patterns of primates, suggesting that new ANN models are needed to more precisely capture the neural mechanisms underlying primate object vision. To this end, large-scale high-resolution behavioral benchmarks, such as those obtained here, could serve as a strong top-down constraint for efficiently discovering such models.

## MATERIALS & METHODS

### Visual images

We examined basic-level, core object recognition behavior using a set of 24 broadly-sampled objects that we previously found to be reliably labeled by independent human subjects, based on the definition of basic-level proposed by (Rosch et al., 1976). For each object, we generated 100 naturalistic synthetic images by first rendering a 3D model of the object with randomly chosen viewing parameters (2D position, 3D rotation and viewing distance), and then placing that foreground object view onto a randomly chosen, natural image background. To do this, each object was first assigned a canonical position (center of gaze), scale (∼2 degrees) and pose, and then its viewing parameters were randomly sampled uniformly from the following ranges for object translation ([-3,3] degrees in both h and v), rotation ([-180,180] degrees in all three axes) and scale ([x0.7, x1.7]. Background images were sampled randomly from a large database of high-dynamic range images of indoor and outdoor scenes obtained from Dosch Design (www.doschdesign.com). This image generation procedure enforces invariant object recognition, rather than image matching, as it requires the visual recognition system (human, animal or model) to tackle the “invariance problem,” the computational crux of object recognition (Ullman and Humphreys, 1996; Pinto et al., 2008). Using this procedure, we previously generated 2400 images (100 images per object) rendered at 1024x1024 pixel resolution with 256-level gray scale and subsequently resized to 256x256 pixel resolution for human psychophysics, monkey psychophysics and model evaluation (Rajalingham et al., 2015). In the current work, we focused our analyses on a randomly subsampled, and then fixed, sub-set of 240 images (10 images per object; here referred to as the “primary test images”). Figure 1A shows the full list of 24 objects, with two example images of each object.

**Figure 1.**
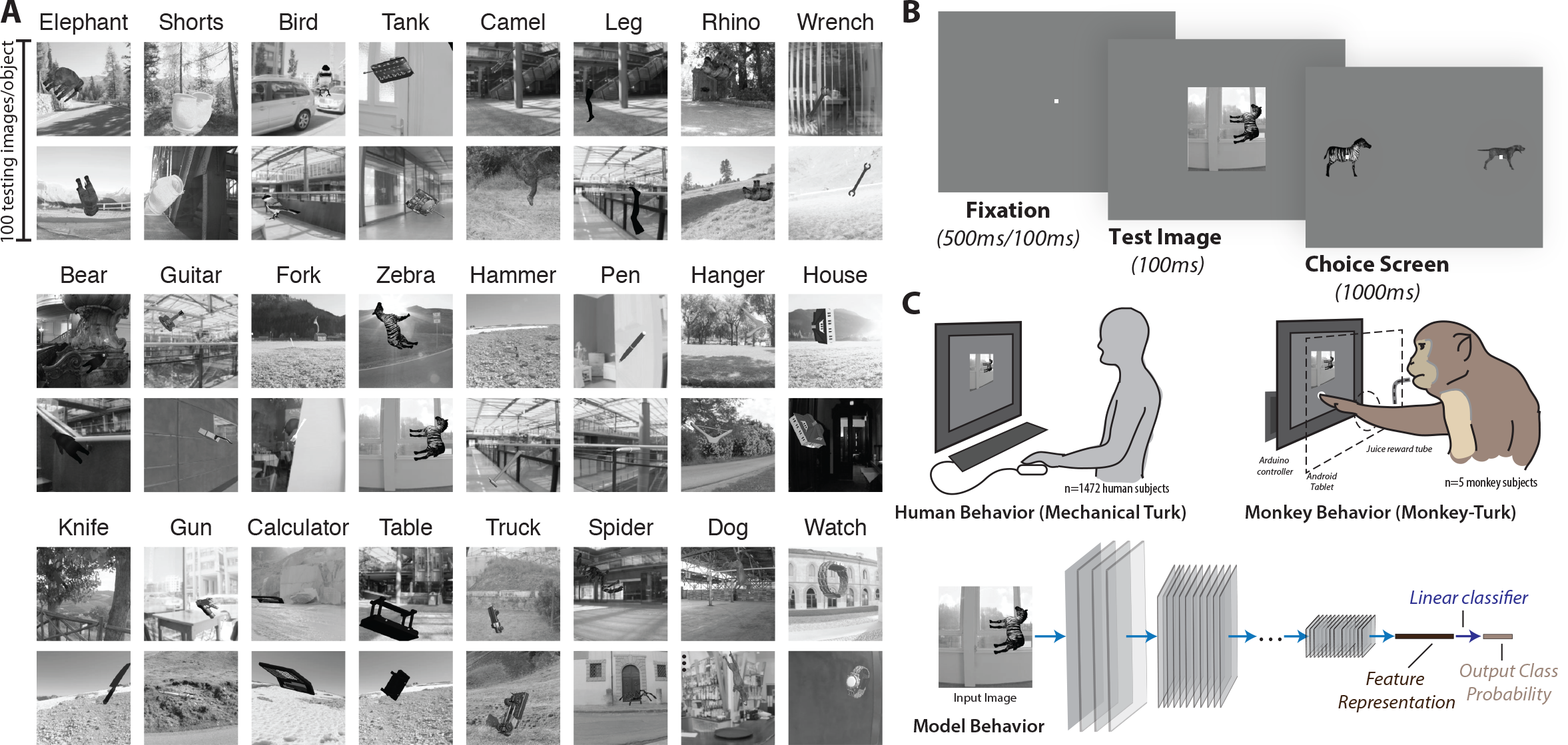
Images and behavioral task. **(A)** Two (out of 100) example images for each of the 24 basic-level objects. To enforce true invariant object recognition behavior, we generated naturalistic synthetic images, each with one foreground object, by rendering a 3D model of each object with randomly chosen viewing parameters and placing that foreground object view onto a randomly chosen, natural image background. **(B)** Time course of example behavioral trial (zebra versus dog) for human psychophysics. Each trial initiated with a central fixation point for 500 ms, followed by 100 ms presentation of a square test image (spanning 6-8° of visual angle). After extinction of the test image, two choice images were shown to the left and right. Human participants were allowed to freely view the response images for up to 1000 ms and responded by clicking on one of the choice images; no feedback was given. To neutralize top-down feature attention, all 276 binary object discrimination tasks were randomly interleaved on a trial-by-trial basis. The monkey task paradigm was nearly identical to the human paradigm, with the exception that trials were initiated by touching a fixation circle horizontally centered on the bottom third of the screen, and successful trials were rewarded with juice while incorrect choices resulted in timeouts of 1–2.5s. **(C)** Large-scale and high-throughput psychophysics in humans (top left), monkeys (top right), and models (bottom). Human behavior was measured using the online Amazon MTurk platform, which enabled the rapid collection ∼1 million behavioral trials from 1472 human subjects. Monkey behavior was measured using a novel custom home-cage behavioral system (MonkeyTurk), which leveraged a web-based behavioral task running on a tablet to test many monkey subjects simultaneously in their home environment. Deep convolutional neural network models were tested on the same images and tasks as those presented to humans and monkeys by extracting features from the penultimate layer of each visual system model and training back-end multi-class logistic regression classifiers. All behavioral predictions of each visual system model were for images that were not seen in any phase of model training.

Because all of the images were generated from synthetic 3D object models, we had explicit knowledge of the viewpoint parameters (position, size, and pose) for each object in each image, as well as perfect segmentation masks. Taking advantage of this feature, we characterized each image based on these high-level attributes, consisting of size, eccentricity, relative pose and contrast of the object in the image. The size and eccentricity of the object in each image were computed directly from the corresponding viewpoint parameters, under the assumption that the entire image would subtend 6° at the center of visual gaze (+/-3° in both azimuth and elevation; see below). For each synthetic object, we first defined its “canonical” 3D pose vector, based on independent human judgments. To compute the relative pose attribute of each image, we estimated the difference between the object’s 3D pose and its canonical 3D pose. Pose differences were computed as distances in unit quaternion representations: the 3D pose (*r_xy_*, *r_xz_*, *r_yz_*) was first converted into unit quaternions, and distances between quaternions *q*_1_, *q*_2_ were estimated as cos^−1^| *q*_1_·*q*_2_ (Huynh, 2009). To compute the object contrast, we measured the absolute difference between the mean of the pixel intensities corresponding to the object and the mean of the background pixel intensities in the vicinity of the object (specifically, within 25 pixels of any object pixel, analogous to computing the local foreground-background luminance difference of a foreground object in an image). Figure 5C shows example images with varying values for the four image attributes.

### Core object recognition behavioral paradigm

Core object discrimination is defined as the ability to discriminate between two or more objects in visual images presented under high view uncertainty in the central visual field (∼10°), for durations that approximate the typical primate, free-viewing fixation duration (∼200 ms) (DiCarlo and Cox, 2007; DiCarlo et al., 2012). As in our previous work (Rajalingham et al., 2015), the behavioral task paradigm consisted of a interleaved set of binary discrimination tasks. Each binary discrimination task is an object discrimination task between a pair of objects (e.g. elephant vs. bear). Each such binary task is balanced in that the test image is equally likely (50%) to be of either of the two objects. On each trial, a test image is presented, followed by a choice screen showing canonical views of the two possible objects (the object that was not displayed in the test image is referred to as the “distractor” object, but note that objects are equally likely to be distractors and targets). Here, 24 objects were tested, which resulted in 276 binary object discrimination tasks. To neutralize feature attention, these 276 tasks are randomly interleaved (trial by trial), and the global task is referred to as a basic-level, core object recognition task paradigm.

### Testing human behavior

All human behavioral data presented here were collected from 1476 human subjects on Amazon Mechanical Turk (MTurk) performing the task paradigm described above. Subjects were instructed to report the identity of the foreground object in each presented image from among the two objects presented on the choice screen (Fig 1B). Because all 276 tasks were interleaved randomly (trial-by-trial), subjects could not deploy feature attentional strategies specific to each object or specific to each binary task to process each test image.

Figure 1B illustrates the time course of each behavioral trial, for a particular object discrimination task (zebra versus dog). Each trial initiated with a central black point for 500 ms, followed by 100 ms presentation of a test image containing one foreground object presented under high variation in viewing parameters and overlaid on a random background, as described above (see *Visual images* above). Immediately after extinction of the test image, two choice images, each displaying a single object in a canonical view with no background, were shown to the left and right. One of these two objects was always the same as the object that generated the test image (i.e., the correct object choice), and the location of the correct object (left or right) was randomly chosen on each trial. After clicking on one of the choice images, the subject was queued with another fixation point before the next test image appeared. No feedback was given; human subjects were never explicitly trained on the tasks. Under assumptions of typical computer ergonomics, we estimate that images were presented at 6–8° of visual angle at the center of gaze, and the choice object images were presented at ±6–8° of eccentricity along the horizontal meridian.

We measured human behavior using the online Amazon MTurk platform (see Figure 1C), which enables efficient collection of large-scale psychophysical data from crowd-sourced “human intelligence tasks” (HITs). The reliability of the online MTurk platform has been validated by comparing results obtained from online and in-lab psychophysical experiments (Majaj et al., 2015; Rajalingham et al., 2015). We pooled 927,296 trials from 1472 human subjects to characterize the aggregate human behavior, which we refer to as the “pooled” human (or “archetypal” human). Each human subject performed only a small number of trials (∼150) on a subset of the images and binary tasks. All 2400 images were used for behavioral testing, but in some of the HITs, we biased the image selection towards the 240 primary test images (1424±70 trials/image on this subsampled set, versus 271±93 trials/image on the remaining images, mean ± SD) to efficiently characterize behavior at image level resolution. Images were randomly drawn such that each human subject was exposed to each image a relatively small number of times (1.5±2.0 trials/image per subject, mean ± SD), in order to mitigate potential alternative behavioral strategies (e.g. “memorization” of images) that could arise from a finite image set. Behavioral signatures at the object-level (B.O1, B.O2, see *Behavioral metrics and signatures)* were measured using all 2400 test images, while image-level behavioral signatures (B.I1n, B.I2n, see *Behavioral metrics and signatures)* were measured using the 240 primary test images. (We observed qualitatively similar results using those metrics on the full 2400 test images, but we here focus on the primary test images as the larger number of trials leads to lower noise levels).

Five other human subjects were separately recruited on MTurk to each perform a large number of trials on the same images and tasks (53,097±15,278 trials/subject, mean ± SD). Behavioral data from these five subjects was not included in the characterization of the pooled human described above, but instead aggregated together to characterize a distinct held-out human pool. For the scope of the current work, this held-out human pool—which largely replicated all behavioral signatures of the larger archetypal human (see Figures 2 and 3)—served as an independent validation of our human behavioral measurements.

**Figure 2.**
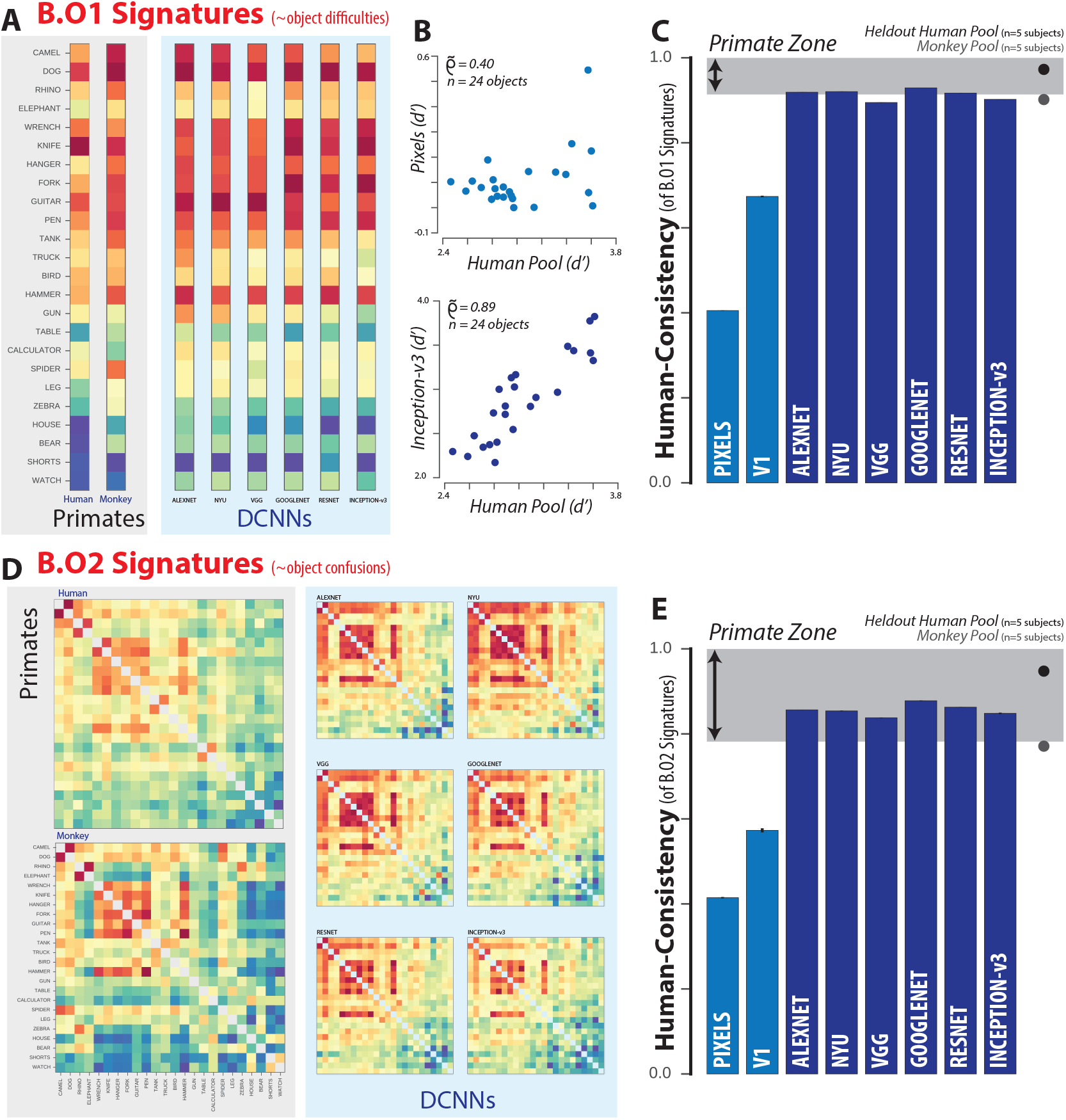
Object-level comparison to human behavior. **(A)** One-versus-all object-level (B.O1) signatures for the pooled human (n=1472 human subjects), pooled monkey (n=5 monkey subjects), and several DCNN_IC_ models. Each B.O1 signature is shown as a 24-dimensional vector using a color scale; each colored bin corresponds to the system’s discriminability of one object against all others that were tested. The color scales span each signature’s full performance range, and warm colors indicate lower discriminability. **(B)** Direct comparison of the B.O1 signatures of a pixel visual system model (top panel) and a DCNN_IC_ visual system model (Inception-v3, bottom panel) against that of the human B.O1 signature. **(C)** *Human-consistency* of B.O1 signatures, for each of the tested model visual systems. The black and gray dots correspond to a held-out pool of five human subjects and a pool of five macaque monkey subjects respectively. The shaded area corresponds to the “primate zone,” a range of consistencies delimited by the estimated *human-consistency* of a pool of infinitely many monkeys (see Figure 4A). **(D)** One-versus-other object-level (B.O2) signatures for pooled human, pooled monkey, and several DCNN_IC_ models. Each B.O2 signature is shown as a 24x24 symmetric matrices using a color scale, where each bin (*i, j*) corresponds to the system’s discriminability of objects *i* and j. Color scales similar to (A). **(E)** Human-consistency of B.O2 signatures for each of the tested model visual systems. Format is identical to (C).

### Testing monkey behavior

Five adult male rhesus macaque monkeys (*Macaca mulatta, subjects M, Z, N, P, B)* were tested on the same basic-level, core object recognition task paradigm described above, with minor modification as described below. All procedures were performed in compliance with National Institutes of Health guidelines and the standards of the Massachusetts Institute of Technology Committee on Animal Care and the American Physiological Society. To efficiently characterize monkey behavior, we used a novel home-cage behavioral system developed in our lab (termed MonkeyTurk, see Fig. 1C). This system leveraged a tablet touchscreen (9” Google Nexus or 10.5” Samsung Galaxy Tab S) and used a web application to wirelessly load the task and collect the data (code available at https://github.com/dicarlolab/mkturk). Analogous to the online Amazon Mechanical Turk, which allows for efficient psychophysical assays of a large number (hundreds) of human users in their native environments, MonkeyTurk allowed us to test many monkey subjects simultaneously in their home environment. Each monkey voluntarily initiated trials, and each readily performed the task a few hours each day that the task apparatus was made available to it. At an average rate of ∼2,000 trials per day per monkey, we collected a total of 836,117 trials from the five monkey subjects over a period of ∼3 months.

Monkey training is described in detail elsewhere (Rajalingham et al., 2015). Briefly, all monkeys were initially trained on the match-test-image-to-object rule using other images and were also trained on discriminating the particular set of 24 objects tested here using a separate set of training images rendered from these objects, in the same manner as the main testing images. Two of the monkeys subjects (Z and M) were previously trained in the lab setting, and the remaining three subjects were trained using MonkeyTurk directly in their home cages and did not have significant prior lab exposure. Once monkeys reached saturation performance on training images, we began the behavioral testing phase to collect behavior on test images. Monkeys did improve throughout the testing phase, exhibiting an increase in performance between the first and second half of trials of 4%±0.9% (mean ± SEM over five monkey subjects). However, the image-level behavioral signatures obtained from the first and the second halves of trials were highly correlated to each other (B.I1 noise-adjusted correlation of 0.85±0.06, mean ± SEM over five monkey subjects, see *Behavioral metrics and signatures* below), suggesting that monkeys did not significantly alter strategies (e.g. did not “memorize” images) throughout the behavioral testing phase.

The monkey task paradigm was nearly identical to the human paradigm (see Figure 1B), with the exception that trials were initiated by touching a white “fixation” circle horizontally centered on the bottom third of the screen (to avoid occluding centrally-presented test images with the hand). This triggered a 100ms central presentation of a test image, followed immediately by the presentation of the two choice images (Fig. 1B, location of correct choice randomly assigned on each trial, identical to the human task). Unlike the main human task, monkeys responded by directly touching the screen at the location of one of the two choice images. Touching the choice image corresponding to the object shown in the test image resulted in the delivery of a drop of juice through a tube positioned at mouth height (but not obstructing view), while touching the distractor choice image resulted in a three second timeout. Because gaze direction typically follows the hand during reaching movements, we assumed that the monkeys were looking at the screen during touch interactions with the fixation or choice targets. In both the lab and in the home cage, we maintained total test image size at ∼6 degrees of visual angle at the center of gaze, and we took advantage of the retina-like display qualities of the tablet by presenting images pixel matched to the display (256 x 256 pixel image displayed using 256 x 256 pixels on the tablet at a distance of 8 inches) to avoid filtering or aliasing effects.

As with Mechanical Turk testing in humans, MonkeyTurk head-free home-cage testing enables efficient collection of reliable, large-scale psychophysical data but it likely does not yet achieve the level of experimental control that is possible in the head-fixed laboratory setting. However, we note that when subjects were engaged in home-cage testing, they reliably had their mouth on the juice tube and their arm positioned through an armhole. These spatial constraints led to a high level of head position trial-by-trial reproducibility during performance of the task paradigm. Furthermore, when subjects were in this position, they could not see other animals as the behavior box was opaque, and subjects performed the task at a rapid pace 40 trials/minute suggesting that they were not frequently distracted or interrupted. The location of the upcoming test image (but not the location of the object within that test image) was perfectly predictable at the start of each behavioral trial, which likely resulted in a reliable, reproduced gaze direction at the moment that each test image was presented. The relatively short—but natural and high performing (Cadieu et al., 2014)—test image duration (100 ms) ensured that saccadic eye movements were unlike to influence test image performance (as they generally take ∼200 ms to initiate in response to the test image, and thus well after the test image has been extinguished).

### Testing model behavior

We tested a number of different deep convolutional neural network (DCNN) models on the exact same images and tasks as those presented to humans and monkeys. Importantly, our core object recognition task paradigm is closely analogous to the large-scale ImageNet 1000-way object categorization task for which these networks were optimized and thus expected to perform well. We focused on publicly available DCNN model architectures that have proven highly successful with respect to this computer vision benchmark over the past five years: AlexNet (Krizhevsky et al., 2012), NYU (Zeiler and Fergus, 2014), VGG (Simonyan and Zisserman, 2014), GoogleNet (Szegedy et al., 2013), Resnet (He et al., 2016), and Inception-v3 (Szegedy et al., 2013). As this is only a subset of possible DCNN models, we refer to these as the DCNN_IC_ (to denote ImageNet-Categorization) visual system model sub-family. For each of the publicly available model architectures, we first used ImageNet-categorization-trained model instances, either using publicly available trained model instances or training them to saturation on the 1000-way classification task in-house. Training took several days on 1-2 GPUs.

We then performed psychophysical experiments on each ImageNet-trained DCNN model to characterize their behavior on the exact same images and tasks as humans and monkeys. We first adapted these ImageNet-trained models to our 24-way object recognition task by re-training the final class probability layer (initially corresponding to the probability output of the 1000-way ImageNet classification task) while holding all other layers fixed. In practice, this was done by extracting features from the penultimate layer of each DCNN_IC_ (i.e. top-most prior to class probability layer), on the same images that were presented to humans and monkeys, and training back-end multi-class logistic regression classifiers to determine the cross-validated output class probability for each image. This procedure is illustrated in Figure 1C. To estimate the hit rate of a given image in a given binary classification task, we renormalized the 24-way class probabilities of that image, considering only the two relevant classes, to sum to one. Object-level and image-level behavioral metrics were computed based on these hit rate estimates (as described in *Behavioral metrics and signatures* below). Importantly, this procedure assumes that the model “retina” layer processes the central 6 degrees of the visual field. It also assumes that linear discriminants (“readouts”) of the model’s top feature layer are its behavioral output (as intended by the model designers). Manipulating either of these choices (e.g. resizing the input images such that they span only part of the input layer, or building linear discriminates for behavior using a different model feature layer) would result in completely new, testable ANN models that we do not test here.

From these analyses, we selected the most *human-consistent* DCNN_IC_ architecture (Inception-v3, see *Behavioral consistency* below), fixed that architecture, and then performed post-hoc analyses in which we varied: the input image sampling, the initial parameter settings prior to training, the filter training images, the type of classifiers used to generate the behavior from the model features, and the classifier training images. To examine input image sampling, we re-trained the Inception-v3 architecture on images from ImageNet that were first spatially filtered to match the spatial sampling of the primate retina (i.e. an approximately exponential decrease in cone density away from the fovea) by effectively simulating a fish-eye transformation on each image. These images were at highest resolution at the “fovea” (i.e. center of the image) with gradual decrease in resolution with increasing eccentricity. To examine the analog of “inter-subject variability”, we constructed multiple trained model instances (“subjects”), where the architecture and training images were held fixed (Inception-v3 and ImageNet, respectively) but the model filter weights initial condition and order of training images were randomly varied for each model instance. Importantly, this procedure is only one possible choice for simulating inter-subject variability for DCNN models, a choice that is an important open research direction that we do not address here. To examine the effect of model training, we fine-tuned an ImageNet-trained Inception-v3 model on a synthetic image set consisting of ∼6.9 million images of 1049 objects (holding out 50,000 images for model validation). These images were generated using the same rendering pipeline as our test images, but the objects were non-overlapping with the 24 test objects presented here. As expected, fine-tuning on synthetic images led to an overall increase in performance of ∼5%. We tested the effect of different classifiers to generate model behavior by testing both multi-class logistic regression and support vector machine classifiers. Additionally, we tested the effect of varying the number of training images used to train those classifiers (20 versus 50 images per class).

### Behavioral metrics and signatures

To characterize the behavior of any visual system, we here introduce four behavioral (B) metrics of increasing richness, requiring increasing amounts of data to measure reliably. Each behavioral metric computes a pattern of unbiased behavioral performance, using a sensitivity index: *d*’ = *Z*(*HitRate*) — *Z*(*FalseAlarmRate*), where Z is the inverse of the cumulative Gaussian distribution. The various metrics differ in the resolution at which hit rates and false alarm rates are computed. Table 1 summarizes the four behavioral metrics, varying the hit-rate resolution (object-level or image-level) and the false-alarm resolution (one-versus-all or one-versus-other). When each metric is applied to the behavioral data of a visual system—biological or artificial—we refer to the result as one behavioral “signature” of that system. Note that we do not consider the signatures obtained here to be the final say on the behavior of these biological or artificial systems—in the terms defined here, new experiments using new objects/images but the same metrics would produce additional behavioral signatures.

**Table 1:** Definition of behavioral performance metrics. The first column provides the name, abbreviation, dimensions, and equations for each of the raw performance metrics. The next two columns provide the definitions for computing the hit rate (HR) and false alarm rate (FAR) respectively.

| Behavioral Metric | Hit Rate | False Alarm Rate |
| --- | --- | --- |
| One-versus all object-level performance<br>(B.O1) ($N_{\text{objects}} \times 1$)<br>$O_1(i) = Z(HR(i)) - Z(FAR(i)),$<br>$i = 1, 2, \dots, N_{\text{objects}}$ | Proportion of trials when images of object $i$ were correctly labeled as object $i$. | Proportion of trials when any image was incorrectly labeled as object $i$. |
| One-versus-other object-level performance<br>B.O2 ($N_{\text{objects}} \times N_{\text{objects}}$)<br>$O_2(i, j) = Z(HR(i, j)) - Z(FAR(i, j)),$<br>$i = 1, 2, \dots, N_{\text{objects}}$<br>$j = 1, 2, \dots, N_{\text{objects}}$ | Proportion of trials when images of object $i$ were correctly labeled as $i$, when presented against distractor object $j$. | Proportion of trials when images of object $j$ were incorrectly labeled as object $i$ |
| One-versus-all image-level performance<br>B.I1 ($N_{\text{images}} \times 1$)<br>$I_1(ii) = Z(HR(ii)) - Z(FAR(ii)),$<br>$ii = 1, 2, \dots, N_{\text{images}}$ | Proportion of trials when image $ii$ was correctly classified as object $i$. | Proportion of trials when any image was incorrectly labeled as object $i$. |
| One-versus-other image-level performance<br>B.I2 ($N_{\text{images}} \times N_{\text{objects}}$)<br>$I_2(ii, j) = Z(HR(ii, j)) - Z(FAR(ii, j)),$<br>$ii = 1, 2, \dots, N_{\text{images}}$<br>$j = 1, 2, \dots, N_{\text{objects}}$ | Proportion of trials when image $ii$ was correctly classified as object $i$, when presented against distractor object $j$. | Proportion of trials when images of object $j$ were incorrectly labeled as object $i$ |

The four behavioral metrics we chose are as follows: First, the one-versus-all object-level performance metric (termed B.O1) estimates the discriminability of each object from all other objects, pooling across all distractor object choices. Since we here tested 24 objects, the resulting B.O1 signature has 24 independent values. Second, the one-versus-other object-level performance metric (termed B.O2) estimates the discriminability of each specific pair of objects, or the pattern of pairwise object confusions. Since we here tested 276 interleaved binary object discrimination tasks, the resulting B.O2 signature has 276 independent values (the off-diagonal elements on one half of the 24x24 symmetric matrix). Third, the one-versus-all image-level performance metric (termed B.I1) estimates the discriminability of each image from all other objects, pooling across all possible distractor choices. Since we here focused on the primary image test set of 240 images (10 per object, see above), the resulting B.I1 signature has 240 independent values. Fourth, the one-versus-other image-level performance metric (termed B.I2) estimates the discriminability of each image from each distractor object. Since we here focused on the primary image test set of 240 images (10 per object, see above) with 23 distractors, the resulting B.I2 signature has 5520 independent values.

Naturally, object-level and image-level behavioral signatures are tightly linked. For example, images of a particularly difficult-to-discriminate object would inherit lower performance values on average as compared to images from a less difficult-to-discriminate object. To isolate the behavioral variance that is specifically driven by image variation and not simply predicted by the objects (and thus already captured by B.O1 and B.O2), we defined normalized image-level behavioral metrics (termed B.I1n, B.I2n) by subtracting the mean performance values over all images of the same object and task. This process is schematically illustrated in Figure 3A. We note that the resulting normalized image-level behavioral signatures capture a significant proportion of the total image-level behavioral variance in our data (e.g. 52%, 58% of human B.I1 and B.I2 variance is driven by image variation, independent of object identity). In this study, we use these normalized metrics for image-level behavioral comparisons between models and primates (see Results).

**Figure 3.**
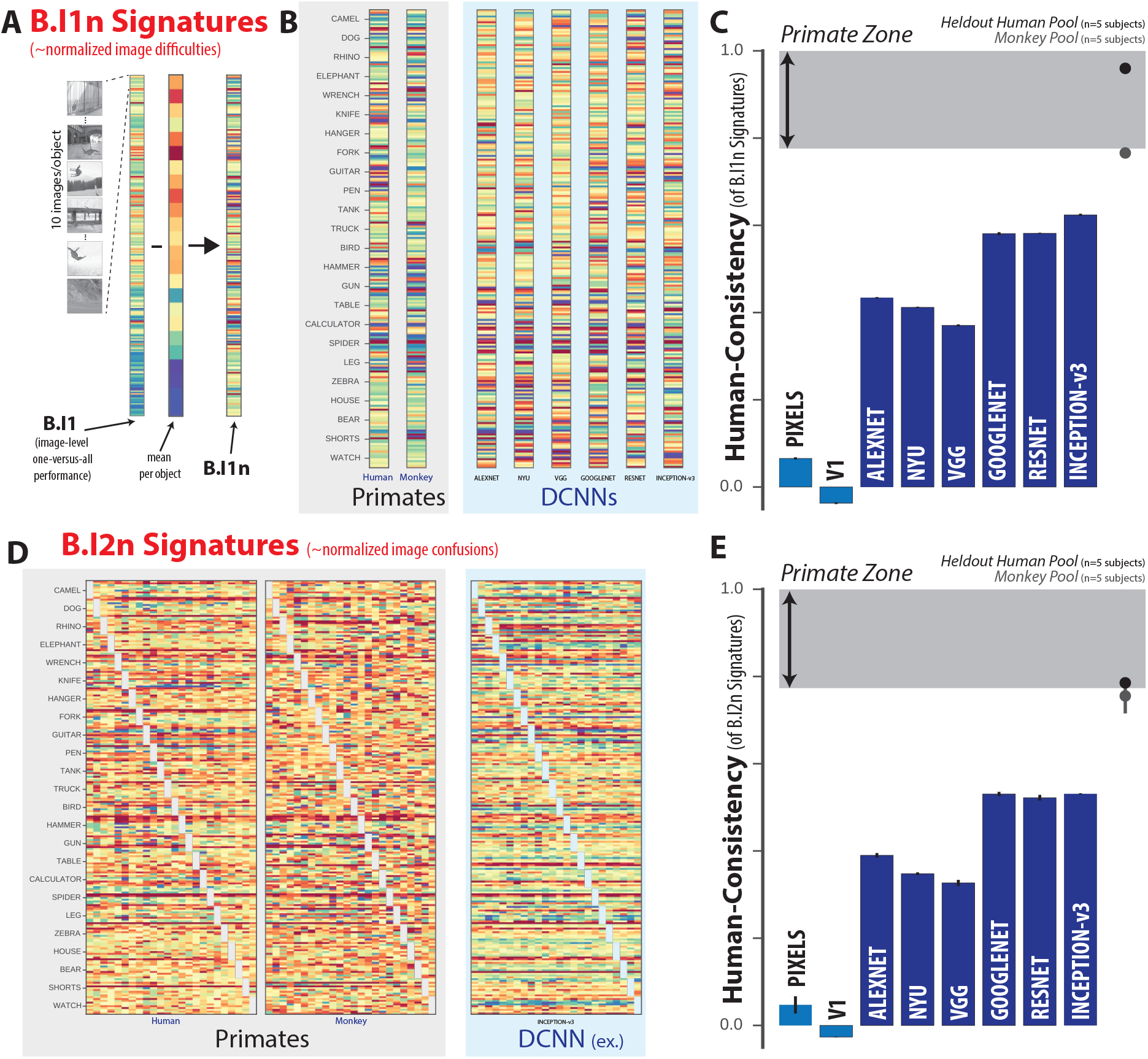
Image-level comparison to human behavior. **(A)** Schematic for computing B.I1n. First, the one-versus-all image-level signature (B.I1) is shown as a 240-dimensional vector (24 objects, 10 images/object) using a color scale, where each colored bin corresponds to the system’s discriminability of one image against all distractor objects. From this pattern, the normalized one-versus-all image-level signature (B.I1n) is estimated by subtracting the mean performance value over all images of the same object. This normalization procedure isolates behavioral variance that is specifically image-driven but not simply predicted by the object. **(B)** Normalized one-versus-all object-level (B.I1n) signatures for the pooled human, pooled monkey, and several DCNN_IC_ models. Each B.I1n signature is shown as a 240-dimensional vector using a color scale, formatted as in (A). Color scales similar to Figure 2A. **(C)** *Human-consistency* of B.Iln signatures for each of the tested model visual systems. Format is identical to Figure 2C. **(D)** Normalized one-versus-other image-level (B.I2n) signatures for pooled human, pooled monkey, and several DCNN_IC_ models. Each B.I2n signature is shown as a 240x24 matrix using a color scale, where each bin (*i, j*) corresponds to the system’s discriminability of image *i* against distractor object *j*. Color scales similar to Figure 2A. **(E)** Human-consistency of B.I2n signatures for each of the tested model visual systems. Format is identical to Figure 2C.

### Behavioral Consistency

To quantify the similarity between a model visual system and the human visual system with respect to a given behavioral metric, we used a measure called the “*human-consistency*’ as previously defined (Johnson et al., 2002). *Human-consistency* (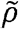) is computed, for each of the four behavioral metrics, as a noise-adjusted correlation of behavioral signatures (DiCarlo and Johnson, 1999). For each visual system, we randomly split all behavioral trials into two equal halves and applied each behavioral metric to each half, resulting in two independent estimates of the system’s behavioral signature with respect to that metric. We took the Pearson correlation between these two estimates of the behavioral signature as a measure of the reliability of that behavioral signature given the amount of data collected, i.e. the split-half internal reliability. To estimate the *human-consistency*, we computed the Pearson correlation over all the independent estimates of the behavioral signature from the model (**m**) and the human (**h**), and we then divide that raw Pearson correlation by the geometric mean of the split-half internal reliability of the same behavioral signature measured for each system: 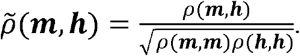

Since all correlations in the numerator and denominator were computed using the same amount of trial data (exactly half of the trial data), we did not need to make use of any prediction formulas (e.g. extrapolation to larger number of trials using Spearman-Brown prediction formula). This procedure was repeated 10 times with different random split-halves of trials. Our rationale for using a reliability-adjusted correlation measure for *human-consistency* was to account for variance in the behavioral signatures that arises from “noise,” i.e., variability that is not replicable by the experimental condition (image and task) and thus that no model can be expected to predict (DiCarlo and Johnson, 1999; Johnson et al., 2002). In sum, if the model (m) is a replica of the archetypal human (h), then its expected human-consistency is 1.0, regardless of the finite amount of data that are collected.

### Characterization of Residuals

In addition to measuring the similarity between the behavioral signatures of primates and models (using *human-consistency* analyses, as described above), we examined the corresponding differences, termed “residual signatures.” Each candidate visual system model’s residual signature was estimated as the residual of a linear least squares regression of the model’s signature on the corresponding human signature and a free intercept parameter. This procedure effectively captures the differences between human and model signatures after accounting for overall performance differences. Residual signatures were estimated on disjoint split-halves of trials, repeating 10 times with random trial permutations. Residuals were computed with respect to the normalized one-versus-all image-level performance metric (B.I1n) as this metric showed a clear difference between DCNN_IC_ models and primates, and the behavioral residual can be interpreted based only the test images (i.e. we can assign a residual per image).

To examine the extent to which the difference between each model and the archetypal human is reliably shared across different models, we measured the Pearson correlation between the residual signatures of pairs of models. Residual similarity was quantified as the proportion of shared variance, defined as the square of the noise-adjusted correlation between residual signatures (the noise-adjustment was done as defined in equation above). Correlations of residual signatures were always computed across distinct split-halves of data, to avoid introducing spurious correlations from subtracting common noise in the human data. We measured the residual similarity between all pairs of tested models, holding both architecture and optimization procedure fixed (between instances of the ImageNet-categorization trained Inception-v3 model, varying in filter initial conditions), varying the architecture while holding the optimization procedure fixed (between all tested ImageNet-categorization trained DCNN architectures), and holding the architecture fixed while varying the optimization procedure (between ImageNet-categorization trained Inception-v3 and synthetic-categorization fine-tuned Inception-v3 models). This analysis addresses not only the reliability of the failure of DCNN_IC_ models to predict human behavior (deviations from humans), but also the relative importance of the characteristics defining similarities within the model sub-family (namely, the architecture and the optimization procedure). We first performed this analysis for residual signatures over the 240 primary test images, and subsequently zoomed in on subsets of images that humans found to be particularly difficult. This image selection was made relative to the distribution of image-level performance of held-out human subjects (B.I1 metric from five subjects); difficult images were defined as ones with performance below the 25^th^ percentile of this distribution.

To examine whether the difference between each model and humans can be explained by simple human-interpretable stimulus attributes, we regressed each DCNN_IC_ model’s residual signature on image attributes (object size, eccentricity, pose, and contrast). Briefly, we constructed a design matrix from the image attributes (using individual attributes, or all attributes), and used multiple linear least squares regression to predict the image-level residual signature. The multiple linear regression was tested using two-fold cross-validation over trials. The relative importance of each attribute (or groups of attributes) was quantified using the proportion of explainable variance (i.e. variance remaining after accounting for noise variance) explained from the residual signature.

### Primate behavior zone

In this work, we are primarily concerned with the behavior of an “archetypal human”, rather than the behavior of any given individual human subject. We operationally defined this concept as the common behavior over many humans, obtained by pooling together trials from a large number of individual human subjects and treating this human pool as if it were acquired from a single behaving agent. Due to inter-subject variability, we do not expect any given human or monkey subject to be perfectly consistent with this archetypal human (i.e. we do not expect it to have a *human-consistency* of 1.0). Given current limitations of monkey psychophysics, we are not yet able to measure the behavior of very large number of monkey subjects at high resolution and consequently cannot directly estimate the *human-consistency* of the corresponding “archetypal monkey” to the human pool. Rather, we indirectly estimated this value by first measuring *human-consistency* as a function of number of individual monkey subjects pooled together (n), and extrapolating the *human-consistency* estimate for pools of very large number of subjects (as n approaches infinity). Extrapolations were done using least squares fitting of an exponential function 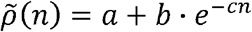 (see Figure 4).

**Figure 4.**
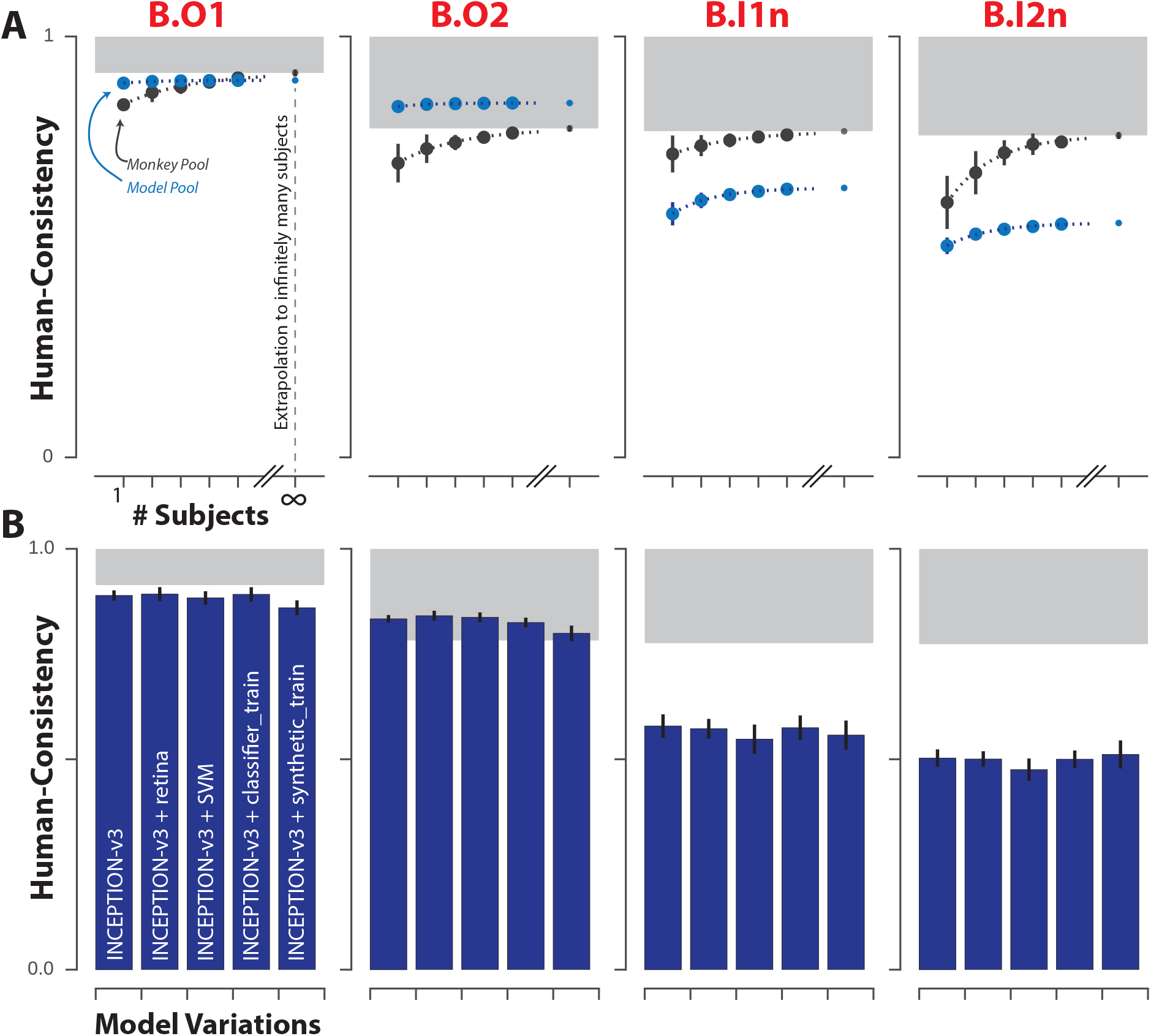
Effect of subject pool size and DCNN model modifications on consistency with human behavior. **(A)** Accounting for natural subject-to-subject variability. For each of the four behavioral metrics, the *human-consistency* distributions of monkey (blue markers) and model (black markers) pools are shown as a function of the number of subjects in the pool (mean ± SD, over subjects). The human consistency increases with growing number of subjects for all visual systems across all behavioral metrics. The dashed lines correspond to fitted exponential functions, and the parameter estimate (mean ± SE) of the asymptotic value, corresponding to the estimated *human-consistency* of a pool of infinitely many subjects, is shown at the right most point on each abscissa. **(B)** Model modifications that aim to rescue the DCNN_IC_ models. We tested several simple modifications (see Methods) to the most *human-consistent* DCNN_IC_ visual system model (Inception-v3). Each panel shows the resulting *human-consistency* per modified model (mean ± SD over different model instances, varying in random filter initializations) for each of the four behavioral metrics.

For each behavioral metric, we defined a “primate zone” as the range of *human-consistency* values delimited by estimates 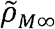 and 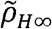 as lower and upper bounds respectively. 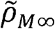 corresponds to the extrapolated estimate of *human-consistency* of a large (i.e. infinitely many) pool of rhesus macaque monkeys; 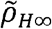 is by definition equal to 1.0. Thus, the primate zone defines a range of *human-consistency* values that correspond to models that accurately capture the behavior of the human pool, at least as well as an extrapolation of our monkey sample. In this work, we defined this range of *human-consistency* values as the criterion for success for computational models of primate visual object recognition behavior.

To make a global statistical inference about whether models sampled from the DCNN_IC_ sub-family meet or fall short of this criterion for success, we attempted to reject the hypothesis that, for a given behavioral metric, the *human-consistency* of DCNN_IC_ models is within the primate zone. To test this hypothesis, we estimated the empirical probability that the distribution of *human-consistency* values, estimated over different model instances within this family, could produce *human-consistency* values within the primate zone. Specifically, we estimated a p-value for each behavioral metric using the following procedure: We first estimated an empirical distribution of Fisher-transformed *human-consistency* values for this model family (i.e. over all tested DCNN_IC_ models and over all trial-resampling of each DCNN_IC_ model). From this empirical distribution, we fit a Gaussian kernel density function, optimizing the bandwidth parameter to minimize the mean squared error to the empirical distribution. This kernel density function was evaluated to compute a p-value, by computing the cumulative probability of observing a *human-consistency* value greater than or equal to the criterion of success (i.e. the Fisher transformed 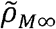 value). This p-value indicates the probability that *human-consistency* values sampled from the observed distribution would fall into the primate zone, with smaller p-values indicating stronger evidence against the hypothesis that the *human-consistency* of DCNN models is within the primate zone.

## RESULTS

In the present work, we systematically compared the basic level core object recognition behavior of primates and state-of-the-art artificial neural network models using a series of behavioral metrics ranging from low to high resolution within a two-alternative forced choice match-to-sample paradigm. The behavior of each visual system, whether biological or artificial, was tested on the same 2400 images (24 objects, 100 images/object) in the same 276 interleaved binary object recognition tasks. Each system’s behavior was characterized at multiple resolutions (see *Behavioral metrics and signatures* in Methods) and directly compared to the corresponding behavioral metric applied on the archetypal human (defined as the average behavior of a large pool of human subjects tested; see Methods). The overarching logic of this study was that, if two visual systems are equivalent, they should produce statistically indistinguishable behavioral signatures with respect to these metrics. Specifically, our goal was to compare the behavioral signatures of visual system models with the corresponding behavioral signatures of primates.

### Object-level behavioral comparison

We first examined the pattern of one-versus-all object-level behavior (termed “B.O1 metric”) computed across all images and possible distractors. Since we tested 24 objects here, the B.O1 signature was 24 dimensional. Figure 2A shows the B.O1 signatures for the pooled human (pooling n=1472 human subjects), pooled monkey (pooling n=5 monkey subjects), and several DCNN_IC_ models as 24-dimensional vectors using a color scale. Each element of the vector corresponds to the system’s discriminability of one object against all others that were tested (i.e. all other 23 objects). The color scales span each signature’s full performance range, and warm colors indicate lower discriminability. For example, red indicates that the tested visual system found the object corresponding to that element of the vector to be very challenging to discriminate from other objects (on average over all 23 discrimination tests, and on average over all images). Figure 2B directly compares the B.O1 signatures computed from the behavioral output of two visual system models—a pixel model (top panel) and a DCNN_IC_ model (Inception-v3, bottom panel)—against that of the human B.O1 signature. We observe a tighter correspondence to the human behavioral signature for the DCNN_IC_ model visual system than for the baseline pixel model visual system. We quantified that similarity using a noise-adjusted correlation between each pair of B.O1 signatures (termed *human-consistency*, following (Johnson et al., 2002)); the noise adjustment means that a visual system that is identical to the human pool will have an expected *human-consistency* score of 1.0, even if it has irreducible trial-by-trial stochasticity; see Methods). Figure 2C shows the B.O1 *human-consistency* for each of the tested model visual systems. We additionally tested the behavior of a held-out pool of five human subjects (black dot) and a pool of five macaque monkey subjects (gray dot), and we observed that both yielded B.O1 signatures that were highly human-consistent (*human-consistency* 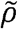 = 0.90, 0.97 for monkey pool and held-out human pool, respectively). We defined a range of *human-consistency* values, termed the “primate zone” (shaded gray area), delimited by extrapolated *human-consistency* estimates of large pools of macaques (see Methods, Figure 4). We found that the baseline pixel visual system model and the low-level V1 visual system model were not within this zone (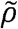 = 0.40, 0.67 for pixels and V1 models, respectively), while all tested DCNN_IC_ visual system models were either within or very close to this zone. Indeed, we could not reject the hypothesis that DCNN_IC_ models are primate-like (p = 0.54, exact test, see Methods).

Next, we compared the behavior of the visual systems at a slightly higher level of resolution. Specifically, instead of pooling over all discrimination tasks for each object, we computed the mean discriminability of each of the 276 pairwise discrimination tasks (still pooling over images within each of those tasks). This yielded a symmetric matrix that is referred to here as the B.O2 signature. Figure 2D shows the B.O2 signatures of the pooled human, pooled monkey, and several DCNN_IC_ visual system models as 24x24 symmetric matrices. Each bin (*i, j*) corresponds to the system’s discriminability of objects *i* and *j*, where warmer colors indicate lower performance; color scales are not shown but span each signature’s full range. We observed strong qualitative similarities between the pairwise object confusion patterns of all of the high level visual systems (e.g. camel and dog are often confused with each other by all three systems). This similarity is quantified in Figure 2E, which shows the *human-consistency* of all examined visual system models with respect to this metric. Similar to the B.O1 metric, we observed that both a pool of macaque monkeys and a held-out pool of humans are highly *human-consistent* with respect to this metric (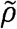 = 0.77, 0.94 for monkeys, humans respectively). Also similar to the B.O1 metric, we found that all DCNN_IC_ visual system models are highly *human-consistent* (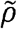 > 0.8) while the baseline pixel visual system model and the low-level V1 visual system model were not (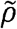 = 0.41, 0.57 for pixels, V1 models respectively). Indeed, all DCNN_IC_ visual system models are within the defined “primate zone” of *human-consistency*, and we could not falsify the hypothesis that DCNN_IC_ models are primate-like (p = 0.99, exact test).

Taken together, humans, monkeys, and current DCNN_IC_ models all share similar patterns of object-level behavioral performances (B.O1 and B.O2 signatures) that are not shared with lower-level visual representations (pixels and V1). However, object-level performance patterns do not capture the fact that some images of an object are more challenging than other images of the same object because of interactions of the variation in the object’s pose and position with the object’s class. To overcome this limitation, we next examined the patterns of behavior at the resolution of individual images on a subsampled set of images where we specifically obtained a large number of behavioral trials to accurately estimate behavioral performance on each image. Note that, from the point of view of the subjects, the behavioral tasks are identical to those already described. We simply aimed to measure and compare their patterns of performance at much higher resolution.

### Image-level behavioral comparison

To isolate purely image-level behavioral variance, i.e. variance that is not predicted by the object and thus already captured by the B.O1 signature, we computed the normalized image-level signature. This normalization procedure is schematically illustrated in Figure 3A which shows that the one-versus-all image-level signature (240-dimensional, 10 images/object) is used to obtain the normalized one-versus-all image-level signature (termed B.I1n, see *Behavioral metrics and signatures)*. Figure 3B shows the B.I1n signatures for the pooled human, pooled monkey, and several DCNN_IC_ models as 240 dimensional vectors. Each bin’s color corresponds to the discriminability of a single image against all distractor options (after subtraction of object-level discriminability, see Figure 3A), where warmer colors indicate lower values; color scales are not shown but span each signature’s full range. Figure 3D shows the *human-consistency* with respect to the B.I1n signature for all tested models. Unlike with object-level behavioral metrics, we now observe a divergence between DCNN_IC_ models and primates. Both the monkey pool and the held-out human pool remain highly *human-consistent* (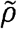 = 0.77, 0.96 for monkeys, humans respectively), but all DCNN_IC_ models were significantly less *human-consistent* (Inception-v3: 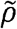 = 0.62) and well outside of the defined “primate zone” of B.I1n *human-consistency*. Indeed, the hypothesis that the *human-consistency* of DCNN_IC_ models is within the primate zone is strongly rejected (p = 6.16e-8, exact test, see Methods).

We can zoom in further by examining not only the overall performance for a given image but also the object confusions for each image, i.e. the additional behavioral variation that is due not only to the test image but to the interaction of that test image with the alternative (incorrect) object choice that is provided after the test image (see Fig. 1B). This is the highest level of behavioral accuracy resolution that our task design allows. In raw form, it corresponds to one-versus-other image-level confusion matrix, where the size of that matrix is the total number of images by the total number of objects (here, 240x24). Each bin (*i,j*) corresponds to the behavioral discriminability of a single image *i* against distractor object *j*. Again, we isolate variance that is not predicted by object-level performance by subtracting the average performance on this binary task (mean over all images) to convert the raw matrix B.I2 above into the normalized matrix, referred to as B.I2n. Figure 3D shows the B.I2n signatures as 240x24 matrices for the pooled human, pooled monkey and top DCNN_IC_ visual system models. Color scales are not shown but span each signature’s full range; warmer colors correspond to images with lower performance in a given binary task, relative to all images of that object in the same task. Figure 3E shows the *human-consistency* with respect to the B.I2n metric for all tested visual system models. Extending our observations using B.I1n, we observe a similar divergence between primates and DCNN_IC_ visual system models on the matrix pattern of image-by-distractor difficulties (B.I2n). Specifically, both the monkey pool and held-out human pool remain highly *human-consistent* (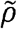 = 0.75, 0.77 for monkeys, humans respectively), while all tested DCNN_IC_ models are significantly less *human-consistent* (Inception-v3: 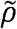 = 0.53) falling well outside of the defined “primate zone” of B.I2n *human-consistency* values. Once again, the hypothesis that the *human-consistency* of DCNN_IC_ models is within the primate zone is strongly rejected (p = 3.17e-18, exact test, see Methods).

### Natural subject-to-subject variation

For each behavioral metric (B.O1, BO2, B.I1n, BI2n), we defined a “primate zone” as the range of consistency values delimited by *human-consistency* estimates 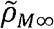 and 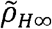 as lower and upper bounds respectively. 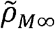 corresponds to the extrapolated estimate of the *human-consistency* of a large (i.e. infinitely many subjects) pool of rhesus macaque monkeys. Thus, the fact that a particular tested visual system model falls outside of the primate zone can be interpreted as a failure of that visual system model to accurately predict the behavior of the archetypal human at least as well as the archetypal monkey.

However, from the above analyses, it is not yet clear whether a visual system model that fails to predict the archetypal human might nonetheless accurately correspond to one or more individual human subjects found within the natural variation of the human population. Given the difficulty of measuring individual subject behavior at the resolution of single images for large numbers of human and monkey subjects, we could not yet directly test this hypothesis. Instead, we examined it indirectly by asking whether an archetypal model—that is a pool that includes an increasing number of model “subjects”—would approach the human pool. We simulated model inter-subject variability by retraining a fixed DCNN architecture with a fixed training image set with random variation in the initial conditions and order of training images. This procedure results in models that can still perform the task but with slightly different learned weight values. We note that this procedure is only one possible choice of generating inter-subject variability within each visual system model type, a choice that is an important open research direction that we do not address here. From this procedure, we constructed multiple trained model instances (“subjects”) for a fixed DCNN architecture, and asked whether an increasingly large pool of model “subjects” better captures the behavior of the human pool, at least as well as a monkey pool. This post-hoc analysis was conducted for the most *human-consistent* DCNN architecture (Inception-v3).

Figure 4A shows, for each of the four behavioral metrics, the measured *human-consistency* of subject pools of varying size (number of subjects n) of rhesus macaque monkeys (black) and ImageNet-trained Inception-v3 models (blue). The *human-consistency* increases with growing number of subjects for both visual systems across all behavioral metrics. To estimate the expected *human-consistency* for a pool of infinitely many monkey or model subjects, we fit an exponential function mapping *n* to the mean *human-consistency* values and obtained a parameter estimate for the asymptotic value (see Methods). We note that estimated asymptotic values are not significantly beyond the range of the measured data—the *human-consistency* of a pool of five monkey subjects reaches within 97% of the *human-consistency* of an estimated infinite pool of monkeys for all metrics—giving credence to the extrapolated *human-consistency* values. This analysis suggests that under this model of inter-subject variability, a pool of Inception-v3 subjects accurately capture archetypal human behavior at the resolution of objects (B.O1, B.O2) by our primate zone criterion (see Figure 4A, first two panels). In contrast, even a large pool of Inception-v3 subjects still fails at its final asymptote to accurately capture human behavior at the image-level (B.I1n, B.I2n) (Figure 4A, last two panels).

### Modification of visual system models to try to rescue their human-consistency

Next, we wondered if some relatively simple changes to the DCNN_IC_ visual system models tested here could bring them into better correspondence with the primate visual system behavior (with respect to B.I1n and B.I2n metrics). Specifically, we considered and tested the following modifications to the most *human-consistent* DCNN_IC_ model visual system (Inception-v3): we (1) changed the input to the model to be more primate-like in its retinal sampling (Inception-v3 + retina-like), (2) changed the transformation (aka “decoder”) from the internal model feature representation into the behavioral output by augmenting the number of decoder training images or changing the decoder type (Inception-v3 + SVM, Inception-v3 + classifier_train), and (3) modified all of the internal filter weights of the model (aka “fine tuning”) by augmenting its ImageNet training with additional images drawn from the same distribution as our test images (Inception-v3 + synthetic-fine-tune). While some of these modifications (e.g. fine-tuning on synthetic images and increasing the number of classifier training images) had the expected effect of increasing mean overall performance (not shown, see Methods), we found that none of these modifications led to a significant improvement in its *human-consistency* on the behavioral metrics (Figure 4B). Thus, the failure of current DCNN_IC_ models to accurately capture the image-level signatures of primates cannot be rescued by simple modifications on a fixed architecture.

### Looking for clues: Image-level comparisons of models and primates

Taken together, Figures 2, 3 and 4 suggest that current DCNN_IC_ visual system models fail to accurately capture the image-level signatures of humans and monkeys. To further examine this failure in the hopes of providing clues for model improvement, we examined the image-level residual signatures of all the visual system models, relative to the pooled human. For each model, we computed its residual signature as the difference (positive or negative) of a linear least squares regression of the model signature on the corresponding human signature. For this analysis, we focused on the B.I1n metric as it showed a clear divergence of DCNN_IC_ models and primates, and the behavioral residual can be interpreted based only on the test images (whereas B.I2n depends on the interaction between test images and distractor choice).

We first asked to what extent the residual signatures are shared between different visual system models. Figure 5A shows the similarity between the residual signatures of all pairs of models; the color of bin (*i,j*) indicates the proportion of explainable variance that is shared between the residual signatures of visual systems *i* and *j*. For ease of interpretation, we ordered visual system models based on their architecture and optimization procedure and partitioned this matrix into four distinct regions. Each region compares the residuals of a “source” model group with fixed architecture and optimization procedure (five Inception-v3 models optimized for categorization on ImageNet, varying only in initial conditions and training image order) to a “target” model group. The target groups of models for each of the four regions are: 1) the pooled monkey, 2) other DCNN_IC_ models from the source group, 3) DCNN_IC_ models that differ in architecture but share the optimization procedure of the source group models and 4) DCNN_IC_ models that differ slightly using an augmented optimization procedure but share the architecture of the source group models. Figure 5B shows the mean (±SD) variance shared in the residuals averaged within these four regions for all images (black dots), as well as for images that humans found to be particularly difficult (gray dots, selected based on held-out human data, see Methods). First, consistent with the results shown in Figure 3, we note that the residual signatures of this particular DCNN_IC_ model are not well shared with the pooled monkey (r^2^=0.39 in region 1), and this phenomenon is more pronounced for the images that humans found most difficult (r^2^=0.17 in region 1). However, this relatively low correlation between model and primate residuals is not indicative of spurious model residuals, as the model residual signatures were highly reliable between different instances of this fixed DCNN_IC_ model, across random training initializations (region 2: r^2^=0.79, 0.77 for all and most difficult images, respectively). Interestingly, residual signatures were still largely shared with other DCNN_IC_ models with vastly different architectures (region 3: r^2^=0.70, 0.65 for all and most difficult images, respectively). However, residual signatures were more strongly altered when the visual training diet of the same architecture was altered (region 4: r^2^=0.57, 0.46 for all and most difficult images respectively, cf. region 3). Taken together, these results indicate that the images where DCNN_IC_ visual system models diverged from humans (and monkeys) were not spurious but were rather highly reliable across different model architectures, demonstrating that current DCNN_IC_ models systematically and similarly diverge from primates.

**Figure 5.**
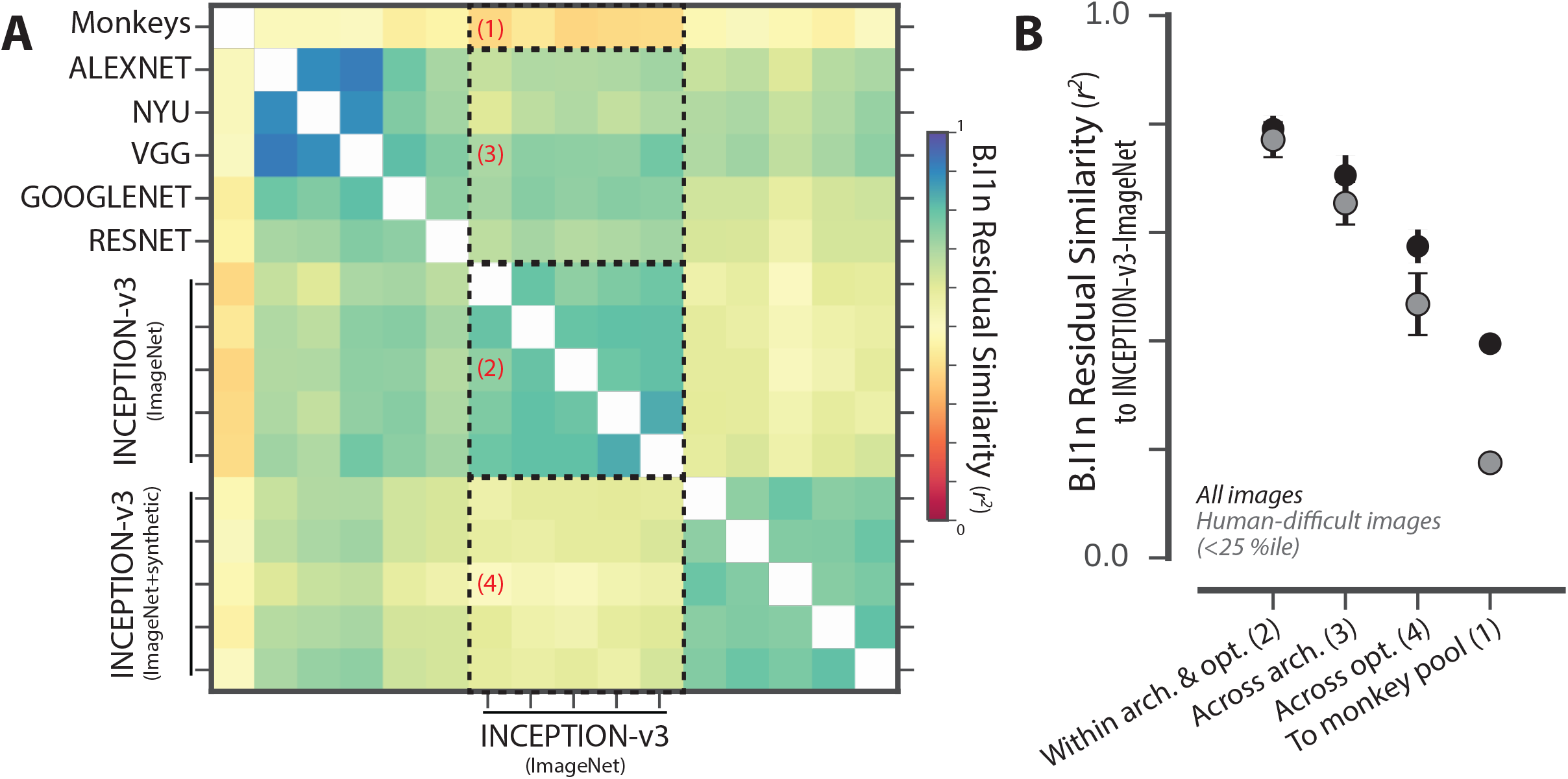
Analysis of unexplained human behavioral variance. **(A)** Residual similarity between all pairs of human visual system models. The color of bin (*i, j*) indicates the proportion of explainable variance that is shared between the residual signatures of visual systems *i* and *j*. For ease of interpretation, we ordered visual system models based on their architecture and optimization procedure and partitioned this matrix into four distinct regions. **(B)** Summary of residual similarity. For each of the four regions in Figure 5A, the similarity to the residuals of Inception-v3 (region 2 in (A)) is shown (mean ± SD, within each region) for all images (black dots), and for images that humans found to be particularly difficult (gray dots, selected based on held-out human data).

To look for clues for model improvement, we asked what, if any, characteristics of images might account for this divergence of models and primates. We regressed the residual signatures of DCNN_IC_ models on four different image attributes (corresponding to the size, eccentricity, pose, and contrast of the object in each image). We used multiple linear regressions to predict the model residual signatures from all of these image attributes, and also considered each attribute individually using simple linear regressions. Figure 6A shows example images (sampled from the full set of 2400 images) with increasing attribute value for each of these four image attributes. While the DCNN_IC_ models were not directly optimized to display primate-like performance dependence on such attributes, we observed that the Inception-v3 visual system model nonetheless exhibited qualitatively similar performance dependencies as primates (see Figure 6B). For example, humans (black), monkeys (gray) and the Inception-v3 model (blue) all performed better, on average, for images in which the object is in the center of gaze (low eccentricity) and large in size. Furthermore, all three systems performed better, on average, for images when the pose of the object was closer to the canonical pose (see Figure 6B); this sensitivity to object pose manifested itself as a non-linear dependence due to the fact that all tested objects exhibited symmetry in at least one axis. The similarity of the patterns in Figure 6B between primates and the DCNN_IC_ visual system models is not perfect but is striking, particularly in light of the fact that these models were not optimized to produce these patterns. However, this similarity is analogous to the similarity in the B.O1 and B.O2 metrics in that it only holds on average over many images. Looking more closely at the image-by-image comparison, we again found that the DCNN_IC_ models failed to capture a large portion of the image-by-image variation (Figure 3). In particular, Figure 6C shows the proportion of variance explained by specific image attributes for the residual signatures of monkeys (black) and DCNN_IC_ models (blue). We found that, taken together, all four of these image attributes explained only ∼10% of the variance in DCNN_IC_ residual signatures, and each individual attribute could explain at most a small amount of residual variance (<5% of the explainable variance). In sum, these analyses show that some behavioral effects that might provide intuitive clues to modify the DCNN_IC_ models are already in place in those models (e.g. a dependence on eccentricity). But the quantitative image-by-image analyses of the remaining unexplained variance (Figure 6C) argue that the DCNN_IC_ visual system models’ failure to capture primate image-level signatures cannot be further accounted for by these simple image attributes and likely stem from other factors.

**Figure 6.**
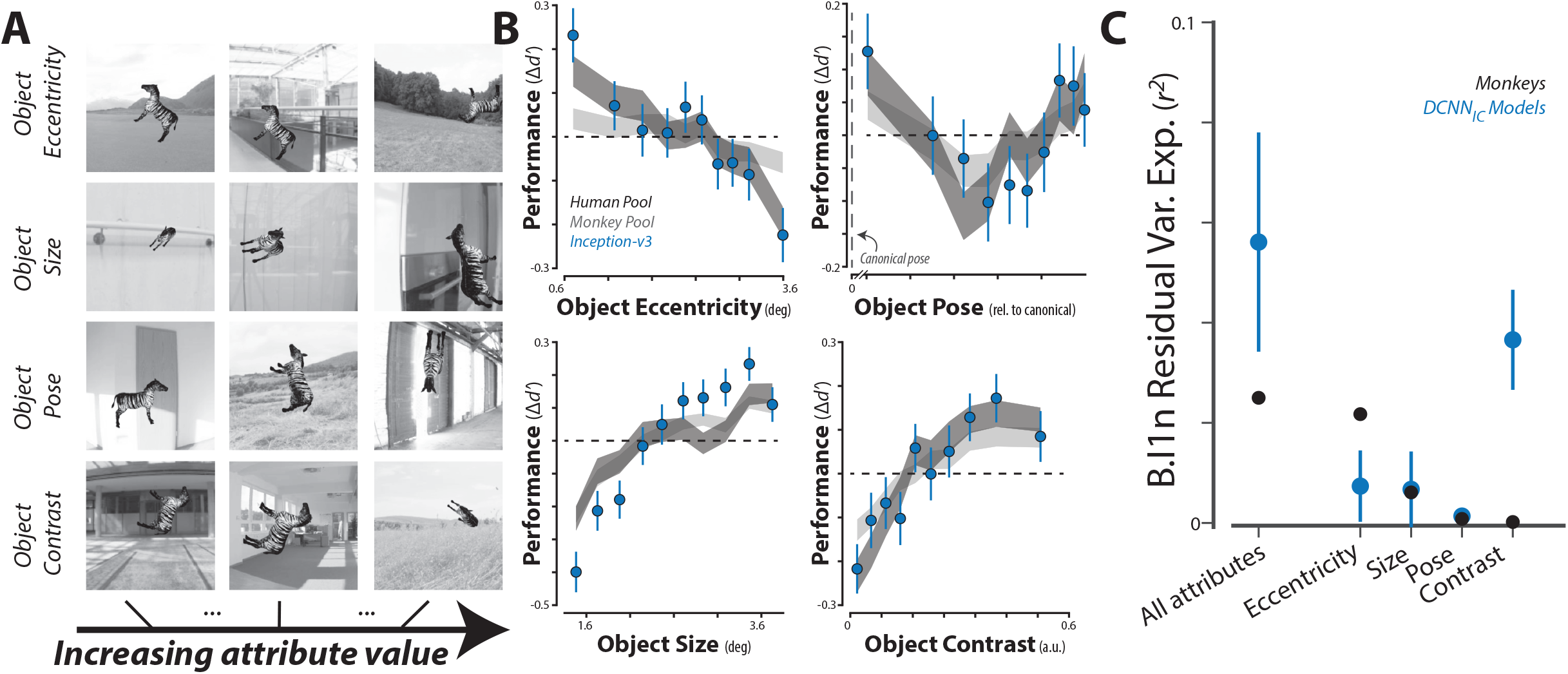
Dependence of primate and DCNN model behavior on image attributes. **(A)** Example images with increasing attribute value, for each of the four pre-defined image attributes (see Methods). **(B)** Dependence of performance (B.I1n) as a function of four image attributes, for humans, monkeys and a DCNN_IC_ model (Inception-v3). **(C)** Proportion of explainable variance of the residual signatures of monkeys (black) and DCNN_IC_ models (blue) that is accounted for by each of the pre-defined image attributes. Error-bars correspond to SD over trial re-sampling for monkeys, and over different models for DCNN_IC_ models.

## DISCUSSION

The current work was motivated by the broad scientific goal of discovering models that quantitatively explain the neuronal mechanisms underlying primate invariant object recognition behavior. To this end, previous work had shown that specific artificial neural network models (ANNs), drawn from a large family of deep convolutional neural networks (DCNNs) and optimized to achieve high levels of object categorization performance on large-scale image-sets, capture a large fraction of the variance in primate visual recognition behaviors (Rajalingham et al., 2015; Jozwik et al., 2016; Kheradpisheh et al., 2016; Kubilius et al., 2016; Peterson et al., 2016; Wallis et al., 2017), and the internal hidden neurons of those same models also predict a large fraction of the image-driven response variance of brain activity at multiple stages of the primate ventral visual stream (Yamins et al., 2013; Cadieu et al., 2014; Khaligh-Razavi and Kriegeskorte, 2014; Yamins et al., 2014; Güçlü and van Gerven, 2015; Cichy et al., 2016; Hong et al., 2016; Seibert et al., 2016; Cadena et al., 2017; Wen et al., 2017). For clarity, we here referred to this sub-family of models as DCNN_IC_ (to denote ImageNet-Categorization training), so as to distinguish them from all possible models in the DCNN family, and more broadly, from the super-family of all ANNs. In this work, we directly compared leading DCNN_IC_ models to primates (humans and monkeys) with respect to their behavioral signatures at both object and image level resolution in the domain of core object recognition. In order to do so, we measured and characterized primate behavior at larger scale and higher resolution than previously possible. We first replicate prior work (Rajalingham et al., 2015) showing that, at the object level, DCNN_IC_ models produce statistically indistinguishable behavior from primates, and we extend that work by showing that these models also match the *average* primate sensitivities to object contrast, eccentricity, size, and pose, a noteworthy similarity in light of the fact that these models were not optimized to produce these performance patterns. However, our primary novel result is that, examining behavior at the higher resolution of individual images, all leading DCNN_IC_ models failed to replicate the image-level behavioral signatures of primates. An important related claim is that rhesus monkeys are more consistent with the archetypal human than any of the tested DCNN_IC_ models (at the image-level).

While it had previously been shown that DCNN_IC_ models can diverge from human behavior on specifically chosen adversarial images (Szegedy et al., 2013), a strength of our work is that we did not optimize images to induce failure but instead randomly sampled the image generative parameter space broadly. As such, our results highlight a *general*, rather than adversarial-induced, failure of DCNN_IC_ models to fully capture the neural mechanisms underlying primate core object recognition behavior. Furthermore, we showed that this failure of current DCNN_IC_ models cannot be explained by simple image attributes and cannot be rescued by simple model modifications (input image sampling, model training, and classifier variations). Taken together, these results suggest that new ANN models are needed to more precisely capture the neural mechanisms underlying primate object vision.

With regards to new ANN models, we can attempt to make prospective inferences about future possible DCNN_IC_ models from the data presented here. Based on the observed distribution of image-level *human-consistency* values for the DCNN_IC_ models tested here, we infer that yet untested model instances sampled identically (i.e. from the DCNN_IC_ model sub-family) are very likely to have similarly inadequate image-level *human-consistency*. While we cannot rule out the possibility that at least one model instance within the DCNN_IC_ sub-family would fully match the image-level behavioral signatures, the probability of sampling such a model is vanishingly small (p<10^−17^ for B.I2n *human-consistency*, estimated using exact test using Gaussian kernel density estimation, see Methods, Results). An important caveat of this inference is that we may have a biased estimate of the *human-consistency* distribution of this model sub-family, as we did not exhaustively sample the sub-family. In particular, if the model sampling process is non-stationary over time (e.g. increases in computational power over time allows larger models to be successfully trained), the *human-consistency* of new (i.e. yet to be sampled) models may lie outside the currently estimated distribution. Consistent with the latter, we observed that current DCNN_IC_ cluster into two distinct “generations” separated in time (before/after the year 2015; e.g. Inception-v3 improves over AlexNet though both lie outside the primate zone in Figure 3). Thus, following this trend, it is possible that the evolution of “next-generation” models within the DCNN_IC_ sub-family could meet our criteria for successful matching primate-like behavior.

Alternatively, it is possible—and we think likely—that future DCNN_IC_ models will also fail to capture primate-like image-level behavior, suggesting that either the architectural limitations (e.g. convolutional, feed-forward) and/or the optimization procedure (including the diet of visual images) that define this model sub-family are fundamentally limiting. Thus, ANN model sub-families utilizing different architectures (e.g. recurrent neural networks) and/or optimized for different behavioral goals (e.g. loss functions other than object classification performance, and/or images other than category-labeled ImageNet images) may be necessary to accurately capture primate behavior. To this end, we propose that testing even individual changes to the DCNN_IC_ models—each creating a new ANN model sub-family—may be the best way forward, because DCNN_IC_ models currently offer the best explanations (in a predictive sense) of both the behavioral and neural phenomena of core object recognition.

To reach that goal of finding a new ANN model sub-family that is a better mechanistic model of the primate ventral visual stream, we propose that even larger-scale, high-resolution behavioral measurements, such as expanded versions of the patterns of image-level performance presented here, could serve as a useful top-down optimization guides. Not only do these high-resolution behavioral signatures have the statistical power to reject the currently leading ANN models, but they can also be efficiently collected at very large scale, in contrast to other guide data (e.g. large-scale neuronal measurements). Indeed, current technological tools for high-throughput psychophysics in humans and monkeys (e.g. Amazon Mechanical Turk for humans, Monkey Turk for rhesus monkeys) enable time- and cost-efficient collection of large-scale behavioral datasets, such as the ∼1 million behavioral trials obtained for the current work. These systems trade off an increase in efficiency with a decrease in experimental control. For example, we did not impose experimental constraints on subjects’ acuity and we can only infer likely head and gaze position. Previous work has shown that patterns of behavioral performance on object recognition tasks from in-lab and online subjects were equally reliable and virtually identical (Majaj et al., 2015), but it is not yet clear to what extent this holds at the resolution of individual images, as one might expect that variance in performance across images is more sensitive to precise head and gaze location. For this reason, we here refrain from making strong inferences from small behavioral differences, such as the small difference between humans and monkeys. Nevertheless, we argue that this sacrifice in exact experimental control while retaining sufficient power for model comparison is a good tradeoff for efficiently collecting large behavioral datasets toward the goal of constraining future models of the primate ventral visual stream.

## ACKNOWLEDGEMENTS

This research was supported by US National Eye Institute grants R01-EY014970 (J.J.D.) and K99-EY022671 (E.B.I.), Office of Naval Research MURI-114407 (J.J.D), IARPA Contract D16PC00002 (J.J.D), Engineering Research Council of Canada (R.R.), and The McGovern Institute for Brain Research

